## Supplementary Information for "Light-sheet photonic force optical coherence elastography for high-throughput quantitative 3D micromechanical imaging"

### Contents

Supplementary Note 1: LS-pfOCE system

Supplementary Note 2: LS-pfOCE reconstruction procedure

Supplementary Note 3: Photothermal response calibration procedure

Supplementary Note 4: Light-sheet radiation-pressure force measurement

Supplementary Note 5: Automated algorithm for depth-resolved force reconstruction based on Radon transform

Supplementary Note 6: Theoretical simulation of light-sheet radiation pressure

Supplementary Data 1: Control experiments for live-cell imaging study

Supplementary Data 2: Normalized stiffness gradient in the pericellular matrix from 3D live-cell micromechanical imaging

Supplementary Data 3: Comparison of cell-modified ECM micromechanical properties in the intermediate regions

Supplementary Data 4: Correlations between changes in micromechanical properties and local matrix deformations during time-lapsed monitoring of Cytochalasin D treatment

Supplementary Discussion 1: Hardware requirements for LS-pfOCE data acquisition

### Supplementary Note 1: LS-pfOCE system

A detailed schematic of the LS-pfOCE system used in the presented experiments is provided in Supplementary Fig. S1. The OCT system uses a broadband superluminescent diode (Thorlabs, LS2000B) as the laser source with a nominal central wavelength and FWHM bandwidth of 1300nm and 200nm, respectively. The fibre-coupled output was launched into a fibre coupler with 50:50 splitting ratio (Thorlabs, TW1300R5A2) which splits into a sample arm and reference arm. In the sample arm, the beam was collimated through a combination of lenses (Thorlabs, $f_2$: AC254-040-C, $f_3$: AC254-050-C and $f_4$: LC1715) and then hit on a two-axis galvo-scanner head (Cambridge Technology, ProSeries 1 10 mm Scan Head with customization to reduce scan angle and improved phase stability). After the beam exits the galvo-scanner, a telescope with 1x magnification ($f_5$: Edmund Optics, 47-318, $f_{11}$: Edmund Optics, 47-317, $f_{12}$: Newport, KPC067; note $f_{11}$ and $f_{12}$ were combined to form a lens pair with an effective focal length around 155 mm) was used to conjugate the galvo-scan mirrors to the back focal plane of the 20X 0.45 NA objective (Olympus, LCPLN20XIR) with a working distance around 8 mm. A protected gold mirror (Thorlabs, PF10-03-M01) mounted on a right-angle kinematic mount (RM4: Thorlabs, KCB1) was used to form an inverted microscope configuration. In the reference arm, the beam was collimated by an achromatic doublet ($f_1$: Thorlabs, AC254-030-C) and reflected by a protected gold mirror, which was mounted on a translatable rail for matching the optical path lengths between sample and reference arms. Interference between backscattered light from the sample and reflected reference beam forms spectral data, which was detected by a spectrometer (Wasatch Photonics, Cobra 1300) with a bandwidth of 245 nm and a 2048-pixel line-scan camera (Sensors Unlimited, GL2048). The spectral data was transferred to a lab computer (Dell, Precision T7610) through a frame grabber (National Instruments, PCIe 1433) and saved as a binary format for further processing. A laser diode (Frankfurt Laser Company, FLU0786M250, HI780 fibre output) with central wavelength of 789 nm was used for PF excitation (pump beam). Fibre-coupled output of the pump beam was collimated by an aspheric lens ($f_6$: Thorlabs, PAFA-X-4-B) and then magnified by a telescope ($f_7$: Thorlabs, AC254-045-B, $f_8$: Thorlabs, AC254-150-B). The light-sheet was generated by a cylindrical lens ($f_9$: Thorlabs, ACY254-100-B). A telescope ($f_{10}$: Thorlabs, AC254-060-B and $f_{11}$, $f_{12}$) was used to further magnify the light-sheet beam and also conjugate the light-sheet to the back focal plane of objective, which was shared with OCT sample arm. A beam steering module (formed by a combination of protected gold mirrors mounted on right-angle kinematic lens mount) was used to align the PF beam to the OCT beam. A dichroic mirror (Thorlabs, DMLP1180R) was used to combine both the OCT sample beam and PF beam. The power of the light-sheet pump beam was harmonically modulated by a laser diode controller (Arroyo Instruments, ComboSource 6340) with a peak power of 120 mW entering the sample. The PF excitation beam modulation signal was synchronized with OCT galvanometer scanning signal via a shared clock on the data acquisition card (National Instruments, PCIe-6353).

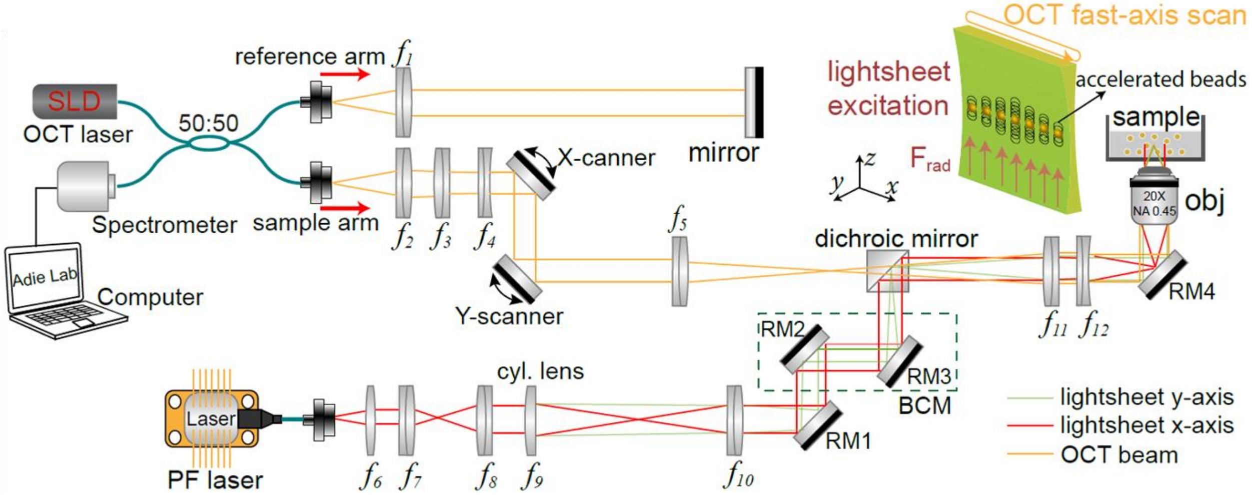


**Supplementary Fig. 1 Detailed schematic of LS-pfOCE system.** The OCT laser is a superluminescent diode (SLD) with wavelength of 1300 ± 100 nm. The PF laser is a fibre-coupled laser diode with wavelength of 789 nm. The long axis (x-axis) of the light-sheet is parallel to fast-axis beam-scanning direction by the X-scanner. RM: right-angle mirror, BCM: beam control module, obj: objective lens.

### Supplementary Note 2: LS-pfOCE reconstruction procedure

The reconstruction of micromechanical properties (i.e., complex shear modulus $G^*$ of medium in the microenvironment of each bead) from raw 3D BM-mode OCT spectral data follows the previously described quantitative procedure[1], with modifications to apply the processing routine on a 3D BM-mode dataset as opposed to an M-mode dataset. A flowchart of the full LS-pfOCE reconstruction procedure is provided in Supplementary Fig. 2. A schematic of the sample configuration and an example cross-sectional OCT image is provided in Supplementary Fig. 3 to illustrate the key components of the image that are relevant to the LS-pfOCE reconstruction.

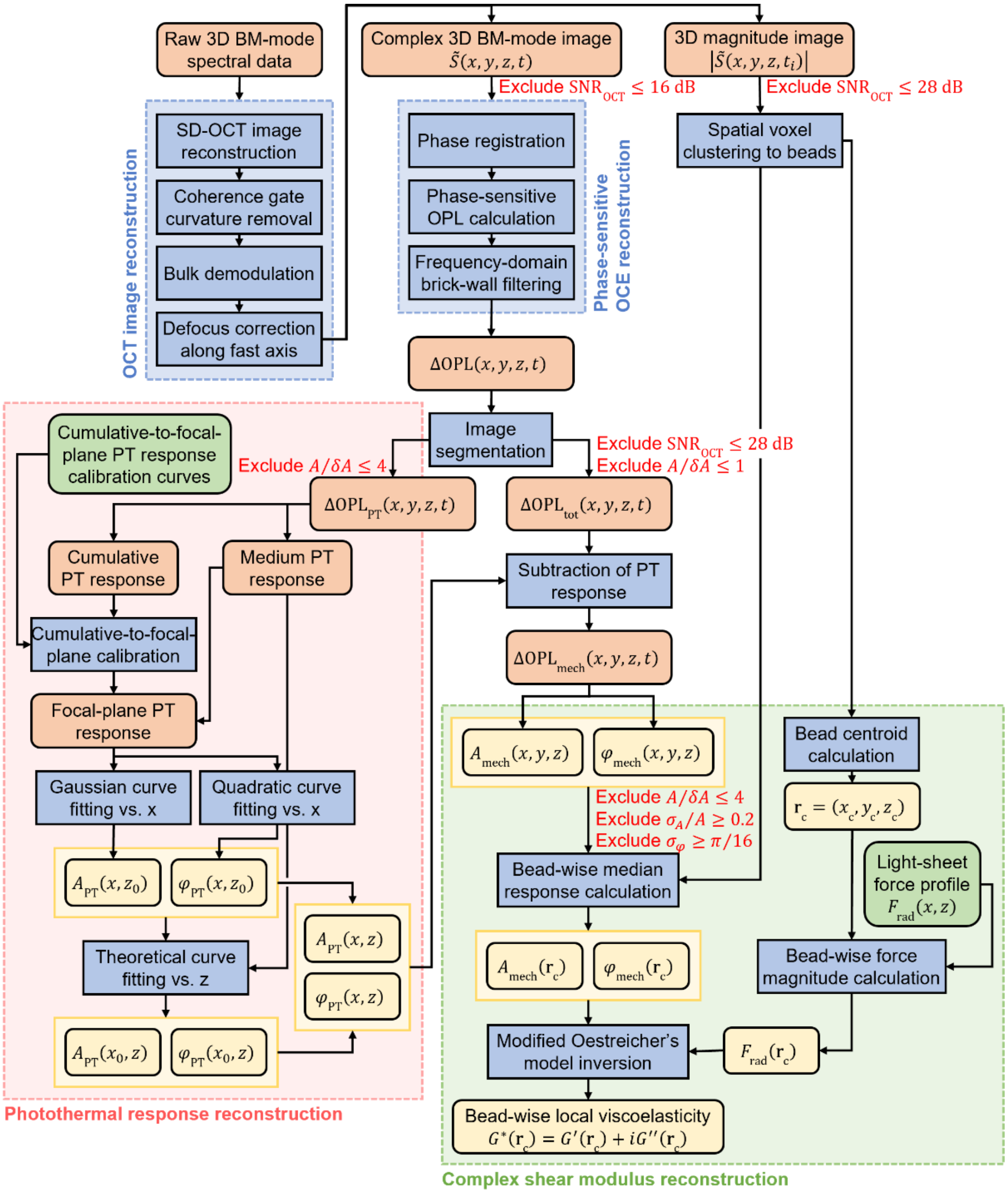


**Supplementary Fig. 2 Data processing flowchart for LS-pfOCE reconstruction.** Red text indicates data exclusion criteria at key processing steps (rationale for these criteria is previously discussed[1]). $\mathrm{SNR}_{OCT}$ : OCT image signal-to-noise ratio, $A/\delta A$: oscillation amplitude-to-noise ratio, $\sigma_A$: standard deviation of mechanical response amplitude over time, $\sigma_\varphi$: standard deviation of mechanical response phase shift over time. See text in this section for definitions of all variables and detailed mathematical descriptions of the processing steps.

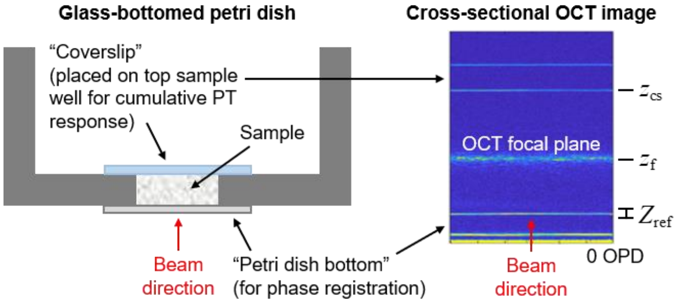


**Supplementary Fig. 3 Schematic of sample configuration and relevant surfaces.** The sample is contained within the well of a glass-bottomed petri dish; whenever applicable, a coverslip is placed on top of the sample well for cumulative photothermal (PT) response measurement. Imaging is performed in an inverted configuration, where the OCT beam and light-sheet enters the sample from the glass-bottom of the petri dish, whose surface is directly in-contact with the sample that is used for phase registration.

**OCT image reconstruction.** Space-domain 3D BM-mode OCT image, $\tilde{S}_{\text{raw}}(x, y, z, t)$, was reconstructed with a standard SD-OCT reconstruction procedure (background subtraction, spectrum resampling, dispersion correction, and inverse Fourier transformation). Then, the image was corrected for defocus along $x$ (fast axis) based on a previously described procedure[2] to obtain a refocused 3D BM-mode image, $\tilde{S}(x, y, z, t)$, for phase-sensitive OCE reconstruction. Briefly, the raw image was flattened via a coherence gate curvature removal procedure using the petri dish bottom surface. Then, the flatted image, $\tilde{S}_{\text{flat}}(x, y, z, t)$, was demodulated to remove the bulk spatial-frequency offset along the fast-axis direction by:

$$\tilde{S}_{\text{demod}}(x, y, z, t) = \tilde{S}_{\text{flat}}(x, y, z, t)\exp(-ik_{x,0}x), \tag{S1}$$

where the bulk modulation along the fast-axis spatial frequency, $k_{x,0}$, was determined as described in Ref.[2]. Lastly, the demodulated image, $\tilde{S}_{\text{demod}}(x, y, z, t)$, was computationally refocused along the fast axis by:

$$\tilde{S}(x, y, z, t) = \mathrm{F}_x^{-1}\left[\mathrm{F}_x\left[\tilde{S}_{\text{demod}}(x, y, z, t)\exp\left(-i(z - z_{\text{f}})\sqrt{(4\pi n/\lambda_0)^2 + k_x^2}\right)\right]\right], \tag{S2}$$

where $\mathrm{F}_x$ and $\mathrm{F}_x^{-1}$ denote the forward and inverse Fourier transform along $x$, respectively. $z_{\text{f}}$, $\lambda_0$, and $n$ denote the axial position of the OCT focal plane, the centre wavelength of the OCT beam, and the refractive index of the medium, respectively. We note that a 2D computational defocus correction along the fast axis was feasible given the BM-mode acquisition scheme, where temporal variation occurred only across the BM-mode frames.

**Phase-sensitive OCE reconstruction.** The OPL response to harmonically-modulated radiation-pressure force, $F_{\text{rad}}$, of each spatial voxel was obtained by extracting the phase of $\tilde{S}(x, y, z, t)$ after phase registration to the petri dish bottom reference surface, as previously describe[1]. Briefly, the axial coordinates of the phase reference surface, $z_{\text{ref}}(x, y)$, was obtained from the magnitude of the OCT image by:

$$z_{\text{ref}}(x, y) = \underset{z \in Z_{\text{ref}}}{\arg\max} \sum_t \left|\tilde{S}(x, y, z, t)\right|, \tag{S3}$$

where $Z_{\mathrm{ref}}$ denotes the depth range containing the petri dish bottom surface for phase registration (Supplementary Fig. 3). Then, the phase-registered OPL response, $\Delta\mathrm{OPL}(x,y,z,t)$, was obtained and expressed as a complex phasor via:

$$\begin{aligned}\Delta\mathrm{OPL}(x,y,z,t) &= \left(\lambda_0/(4\pi n)\right)\angle\left[\tilde{S}(x,y,z,t)\tilde{S}^*(x,y,z,t)\right] \\ &= A(x,y,z)\exp\left(i\varphi(x,y,z)\right)\tilde{h}(t)\end{aligned}, \quad \text{(S4)}$$

where $S^*$ represents the complex conjugate of $S$. The phasor notation expresses the complex OPL response by its amplitude $A(x,y,z)$, phase shift w.r.t. the drive waveform $\varphi(x,y,z)$, and the complex drive waveform $\tilde{h}(t)=\exp(i(\omega t-\pi))$, where $\omega = 2\pi\times 20\ \mathrm{Hz}$ denotes the angular modulation frequency of the harmonically-modulated $F_{\mathrm{rad}}$. Subsequent mention of "phase shift" shall refer to the phase shift w.r.t. to the drive waveform as defined in Eq. (S4). Lastly, the induced response at 20 Hz was extracted from $\Delta\mathrm{OPL}(x,y,z,t)$ via a narrow-band brick-wall filter (passband 20 ± 0.2 Hz) in the frequency domain.

**Image segmentation.** The $\Delta\mathrm{OPL}(x,y,z,t)$ was divided into the photothermal (PT) and total response data regions based on the time-averaged magnitude of $\tilde{S}(x,y,z,t)$ at each spatial voxel[3]. The threshold values used for PAAm gels with 1.7-µm polystyrene probe beads are: $600\le|\tilde{S}|<4000$ and $|\tilde{S}|\ge 4200$ for PT and total response data regions, respectively. The threshold values used for collagen matrices and live-cell imaging study with 1.9-µm melamine-resin probe beads are: $600\le|\tilde{S}|<10000$ and $|\tilde{S}|\ge 10200$ for PT and total response data regions, respectively. In addition, spatial voxels within a 6-µm radius from the surface of each probe beads were excluded from the PT data region, under the premise that local deformation due to the oscillation of the beads extend into the medium in the vicinity of each bead. Lastly, the exclusion criteria indicated in Supplementary Fig. 2 were applied as previously described[1].

**Photothermal response reconstruction.** The PT response of the medium in LS-pfOCE varies as a function of both $x$ and $z$ due to the Gaussian intensity profile of the light-sheet along its long axis. Thus, the lateral profile of the PT response must be reconstructed in addition to the depth-dependent PT response previously described[1,3]. Another distinction in the LS-pfOCE measurements presented here is that no exogenous PT reporters (e.g., 0.1-µm polystyrene beads in Ref.[1,3]) were added to the sample. Only the endogenous scattering signal from the medium and the cumulative PT response at the coverslip (placed on top of the sample well, Supplementary Fig. 3) were used to reconstruct the PT response. This modification was made to minimize the amount of exogenous material added to the sample, particularly for the purpose of the live-cell imaging study.

The PT response reconstruction for LS-pfOCE can be divided into two steps: 1) reconstruction of PT response amplitude and phase shift as a function of $x$ at the focal plane of the OCT beam (where OCT scattering signal is strongest), $A_{\mathrm{PT}}(x,z_{\mathrm{f}})$ and $\varphi_{\mathrm{PT}}(x,z_{\mathrm{f}})$, and 2) reconstruction of depth-dependent PT response amplitude and phase shift at the lateral centre of the light-sheet $A_{\mathrm{PT}}(x_0,z)$ and $\varphi_{\mathrm{PT}}(x_0,z)$. In the absence of exogenous PT reporters, the first step relies on measurements of PT response from the endogenous scattering signal of the medium at the depth $z_{\mathrm{f}}$, which are limited due to the weak scattering of the hydrogel medium. Thus, the direct measurements of PT response at the focal plane were combined with the cumulative PT response at the coverslip $A_{\mathrm{PT}}(x,z_{\mathrm{cs}})$ and $\varphi_{\mathrm{PT}}(x,z_{\mathrm{cs}})$, after a cumulative-to-focal plane calibration procedure (see Supplementary Note 3 for details). The resulting $A_{\mathrm{PT}}(x,z_{\mathrm{f}})$ and $\varphi_{\mathrm{PT}}(x,z_{\mathrm{f}})$ data were fit to a Gaussian and a quadratic curve as a function of $x$, respectively.

Then, the second step compiles all available measurements (i.e., both the direct measurements and the calibrated response) at lateral coordinate $x_0$ to perform the depth-dependent PT response curve fitting[1]. Lastly, the full 2D PT response amplitude and phase as a function of $x$ and $z$ were obtained from:

$$A_{\mathrm{PT}}(x,z) = A_{\mathrm{PT}}(x_0,z)\left[A_{\mathrm{PT}}(x,z_{\mathrm{f}})/A_{\mathrm{PT}}(x_0,z_{\mathrm{f}})\right], \tag{S5}$$

$$\varphi_{\mathrm{PT}}(x,z) = \varphi_{\mathrm{PT}}(x_0,z) + \left[\varphi_{\mathrm{PT}}(x,z_{\mathrm{f}}) - \varphi_{\mathrm{PT}}(x_0,z_{\mathrm{f}})\right]. \tag{S6}$$

An example of the full 2D PT response reconstructed in PAAm is shown in Supplementary Fig. 4.

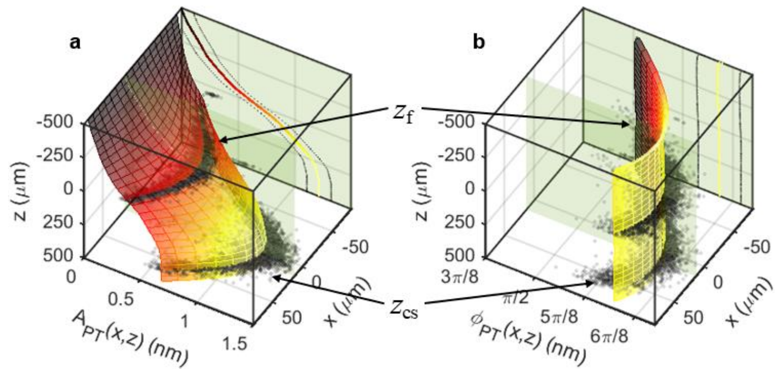


**Supplementary Fig. 4 Reconstructed 2D profiles of LS-pfOCE photothermal response. a**, PT response amplitude and **b**, phase of PAAm gel. Black markers represent measurements of focal-plane and cumulative (coverslip) PT responses. Surface plots represent the reconstructed 2D responses from Eqs. (S5) and (S6). Depth-dependent profiles at the light-sheet lateral centre (green plane) are also shown.

**Isolation of bead mechanical response.** Mechanical response at each spatial voxel was obtained by subtracting the reconstructed PT response from the measured total response, i.e., $\Delta\mathrm{OPL}(x,y,z,t)$ in the total response data region, and expressed as a complex phasor via:

$$\begin{aligned}\Delta\mathrm{OPL}_{\mathrm{mech}}(x,y,z,t) &= \Delta\mathrm{OPL}_{\mathrm{tot}}(x,y,z,t) - A_{\mathrm{PT}}(x,z)\exp\left(i\varphi_{\mathrm{PT}}(x,z)\right)\tilde{h}(t) \\ &= A_{\mathrm{mech}}(x,y,z)\exp\left(i\varphi_{\mathrm{mech}}(x,y,z)\right)\tilde{h}(t)\end{aligned}, \tag{S7}$$

where $A_{\mathrm{mech}}(x,y,z)$ and $\varphi_{\mathrm{mech}}(x,y,z)$ are the mechanical response amplitude and phase shift at each spatial voxel, respectively. Then, the exclusion criteria indicated in Supplementary Fig. 2 were applied as previously described[1]. To obtain the mechanical response of each probe bead, the available spatial voxels were clustered into individual beads based on their spatial coordinates using a density-based spatial clustering algorithm (MATLAB "dbscan" function). Then, the bead-wise mechanical response amplitude and phase were obtained from the median of all voxels that made up each bead by:

$$A_{\mathrm{mech}}(\mathbf{r}_{\mathrm{c}}) = \mathrm{median}\left[A_{\mathrm{mech}}(x,y,z)\right]\Big|_{x\in X_{\mathrm{b}},y\in Y_{\mathrm{b}},z\in Z_{\mathrm{b}}}, \tag{S8}$$

$$\varphi_{\mathrm{mech}}(\mathbf{r}_{\mathrm{c}}) = \mathrm{median}\left[\varphi_{\mathrm{mech}}(x,y,z)\right]\Big|_{x\in X_{\mathrm{b}},y\in Y_{\mathrm{b}},z\in Z_{\mathrm{b}}}, \tag{S9}$$

where $X_{\mathrm{b}}$, $Y_{\mathrm{b}}$, $Z_{\mathrm{b}}$ represent vectors $x$, $y$, and $z$ coordinates of the spatial voxels that make up each bead and $\mathbf{r}_{\mathrm{c}}$ denotes the position vector of the bead centroid.

**Complex shear modulus reconstruction.** The complex shear modulus of the medium in the vicinity of each bead, $G^*(\mathbf{r}_{\mathrm{c}})$, was reconstructed from the bead-wise mechanical response, $A_{\mathrm{mech}}(\mathbf{r}_{\mathrm{c}})$ and $\varphi_{\mathrm{mech}}(\mathbf{r}_{\mathrm{c}})$, and the force magnitude at the bead centroid, $F_{\mathrm{rad}}(\mathbf{r}_{\mathrm{c}})$ (see Supplementary Note 3 for details). We note that $A_{\mathrm{mech}}$, $\varphi_{\mathrm{mech}}$, and $G^*$ are all a function of $\omega$ (i.e., frequency-dependent), but the argument $\omega$ will be omitted in the following derivation for simplicity (given that $\omega$ was kept constant for all results presented in this manuscript). We will only specify the argument $\mathbf{r}_c$ to emphasize the reconstruction of the micromechanical properties of the medium at the location of each bead. The frequency-domain equation of motion of a sphere undergoing oscillatory motion in a viscoelastic medium by an external harmonically-modulated $F_{rad}$ is given by[5]:

$$-m\omega^2 A_{mech}(\mathbf{r}_c)\exp(i\varphi_{mech}(\mathbf{r}_c))\tilde{h}(t) = F_{rad}(\mathbf{r}_c)\tilde{h}(t) - 6\pi a\tilde{G}_{eff}(\mathbf{r}_c)A_{mech}(\mathbf{r}_c)\exp(i\varphi_{mech}(\mathbf{r}_c))\tilde{h}(t), \qquad (S10)$$

where $m$ and $a$ denote mass and radius of the oscillating sphere, respectively. The LHS in Eq. (S10) corresponds to the inertial term, which can be negligible for small $m$ and $\omega$. The first and second terms on the RHS corresponds to the externally applied force and the impedance of the oscillating sphere due to the surrounding viscoelastic medium, respectively. The "effective modulus" $\tilde{G}_{eff}$ for an incompressible linear viscoelastic medium is given as a function of $G^*$ according to Oestreicher's model of impedance of an oscillating sphere in a viscoelastic medium[4]:

$$\tilde{G}_{eff}(\mathbf{r}_c) = G^*(\mathbf{r}_c)\left[1 - ik^*(\mathbf{r}_c)a - \tfrac{1}{9}k^{*2}(\mathbf{r}_c)a^2\right], \qquad (S11)$$

where $k^*$ denotes the complex shear wave number in the viscoelastic medium. On the RHS of Eq. (S11), the $ik^*a$ term accounts for the damping due to the radiation of the shear wave caused by the sphere oscillation while the $\frac{1}{9}k^{*2}a^2$ term accounts for the induced mass of the medium that oscillates with the sphere[5]. (We note that Eqs. (S10) and (S11) would reduce to the Generalized Stokes-Einstein Relation commonly used in microrheology if these two terms and the inertial term in Eq. (S10) were negligible.)

The micromechanical properties of the medium at each bead location were obtained by first solving Eq. (S10) for $\tilde{G}_{eff}(\mathbf{r}_c)$ using the experimentally measured $A_{mech}(\mathbf{r}_c)$, $\varphi_{mech}(\mathbf{r}_c)$, and $F_{rad}(\mathbf{r}_c)$. Then, Eq. (S11) was inverted to obtain $G^*(\mathbf{r}_c)$ via:

$$G^*(\mathbf{r}_c) = \frac{\rho\omega^2}{k^{*2}(\mathbf{r}_c)} = \frac{4\rho\omega^2\tilde{k}_{eff}^{\,2}(\mathbf{r}_c)}{\sqrt{4\tilde{k}_{eff}(\mathbf{r}_c) - a^2} - ia}, \qquad (S12)$$

where $\rho$ denotes the mass density of the medium and $\tilde{k}_{eff}$ was computed from $\tilde{G}_{eff}$ by:

$$\tilde{k}_{eff}(\mathbf{r}_c) = a^2/9 + \tilde{G}_{eff}(\mathbf{r}_c)/(\rho\omega^2). \qquad (S13)$$

### Supplementary Note 3: Photothermal response calibration procedure

The main challenge in the reconstruction of PT response for LS-pfOCE compared to the previously reported Gaussian-beam PF-OCE[1,3] is the need to also characterize the lateral profile of the light-sheet PT response in addition to the depth-dependent (axial) profile. This task requires independent measurements of the PT response inside the medium (at the OCT focal plane, about which LS-pfOCE measurements are taken) as a function of $x$, which is made even more challenging in the absence of exogenous PT reporters. The general principle to mitigate this challenge was to implement a PT calibration procedure, where the relationship between the focal-plane PT response inside the medium, $A_{\mathrm{PT}}(x, z_{\mathrm{f}})$ and $\varphi_{\mathrm{PT}}(x, z_{\mathrm{f}})$, and the cumulative PT response at the coverslip (see Supplementary Fig. 3 for schematic), $A_{\mathrm{PT}}(x, z_{\mathrm{cs}})$ and $\varphi_{\mathrm{PT}}(x, z_{\mathrm{cs}})$, was pre-characterized in a "PT calibration sample" using the same LS-pfOCE acquisition protocol. The "PT calibration sample" provided independent measurements of both the focal-plane and the cumulative PT responses, which were used to compute the cumulative-to-focal-plane calibrations curves for the PT response amplitude, $\varepsilon_A(x)$, and phase, $\varepsilon_\varphi(x)$, via:

$$\varepsilon_A(x) = A_{\mathrm{PT}}(x, z_{\mathrm{f}}) / A_{\mathrm{PT}}(x, z_{\mathrm{cs}}), \tag{S14}$$

$$\varepsilon_\varphi(x) = \varphi_{\mathrm{PT}}(x, z_{\mathrm{cs}}) - \varphi_{\mathrm{PT}}(x, z_{\mathrm{f}}). \tag{S15}$$

After characterizing $\varepsilon_A(x)$ and $\varepsilon_\varphi(x)$, $A_{\mathrm{PT}}(x, z_{\mathrm{f}})$ and $\varphi_{\mathrm{PT}}(x, z_{\mathrm{f}})$ could be estimated from $A_{\mathrm{PT}}(x, z_{\mathrm{cs}})$ and $\varphi_{\mathrm{PT}}(x, z_{\mathrm{cs}})$ measured at the coverslip in each sample using Eqs. (S14) and (S15) in the subsequent LS-pfOCE experiments. These "calibrated" focal-plane PT responses were combined with any available (albeit limited) direct measurements from the intrinsic OCT scattering signal at the focal plane to reconstruct (via Gaussian fit) lateral profile of the PT response in each sample. The specific PT calibration procedure and the "PT calibration sample" differ for each type of samples, depending on the sample characteristics and the specific requirements for each experiment, as outlined below for PAAm gels, collagen matrices, and cell-seeded fibrin constructs for live-cell imaging. This also implies that the procedures outlined below may be modified and tailored to different types of samples and experiments in future applications of LS-pfOCE.

**PAAm gels.** The PAAm gels appeared almost entirely transparent on the OCT image in the absence of exogenous PT reporters. Thus, the "PT calibration sample" in this case was a PAAm gel with high stiffness (6T1C PAAm with $G' > 2$ kPa, see Table 1 in Ref.[1]) containing 0.5-µm diameter polystyrene beads (14-µm average bead spacing). These 0.5-µm polystyrene beads served as the PT reporters under the premise that the low peak light-sheet $F_{\mathrm{rad}}$ magnitude exerted on these beads (order of 0.1 pN) produce negligible (i.e., below the supported displacement sensitivity of the system) mechanical response (order of 10 pm) in the high-stiffness PAAm gel. This was similar to the differential scattering approach implemented in Ref.[3] with 0.1-µm polystyrene beads as PT reporters. However, the higher OCT scattering intensity of the 0.5-µm beads used here provided a more reliable independent measurements of $\Delta\mathrm{OPL}$ at the focal plane owing to the OCT SNR-limited displacement sensitivity of phase-sensitive OCT[3]. The $\varepsilon_A(x)$ and $\varepsilon_\varphi(x)$ calibration curves were obtained from independent measurements of cumulative (from the coverslip) and focal-plane (from the 0.5-µm beads at depth $z_{\mathrm{f}}$) PT responses (Supplementary Fig. 5a, b). Lastly, we verified that the relative depth-dependent profile of both $A_{\mathrm{PT}}$ and $\varphi_{\mathrm{PT}}$ in the softer 3T1C and 3T2C PAAm gels (used in Figs. 1e and 2a, b, d) were consistent with that of the stiffer 6T1C PT calibration sample, using measurements from Gaussian-beam PF-OCE[1] (Supplementary Fig. 5c).

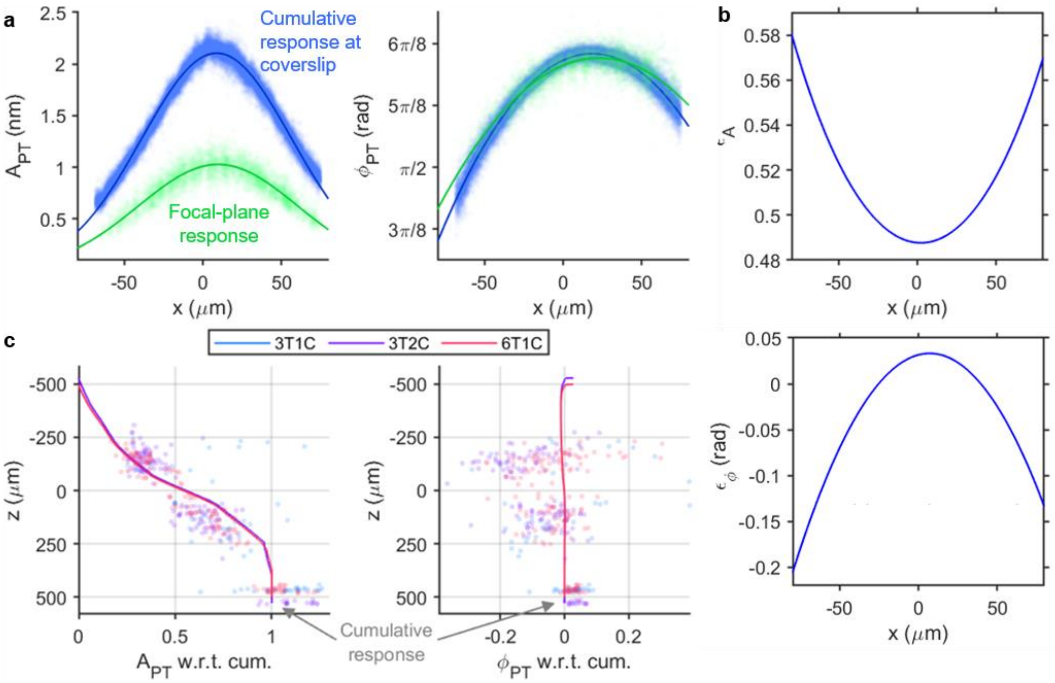


**Supplementary Fig. 5 Cumulative-to-focal-plane PT response calibration in PAAm gels.** **a**, Cumulative (blue) and focal-plane (green) PT response amplitude (left) and phase (right) measured in 6T1C PAAm gel "PT calibration sample". Markers represent measurements. Curves represent Gaussian and quadratic fit for $A_{\mathrm{PT}}$ and $\varphi_{\mathrm{PT}}$, respectively. **b**, Cumulative-to-focal plane calibration curves $\varepsilon_A(x)$ (top) and $\varepsilon_\varphi(x)$ (bottom) obtained from measurements in **a**. **c**, Relative depth-dependent $A_{\mathrm{PT}}$ (left) and $\varphi_{\mathrm{PT}}$ (right) in 3T1C, 3T2C, and 6T1C PAAm gels, each normalized to its cumulative response value at the coverslip. Markers represent measurements. Curves represent depth-dependent curve fit as described in Ref.[1]. The relative depth-dependent profiles (i.e., cumulative-to-focal-plane relationship) are consistent in all PAAm concentrations.

**Collagen matrices.** The fibrous collagen matrices generated intrinsic OCT scattering signal from the collagen fibres, but the available PT response measurements at the focal plane (after the exclusion criteria and excluding any voxels within a 6-µm radius of each bead, see Supplementary Note 1) alone were still insufficient for a reliable Gaussian fit to obtain the lateral profile of the focal-plane PT response. Thus, direct measurements at the focal plane still needed to be supplemented by the cumulative-to-focal-plane calibration procedure. In this case, the "PT calibration sample" was simply another replicate of the same collagen matrix but containing no probe beads (1.9-µm melamine-resin beads), where all available OCT scattering signal at the focal plane that passed the exclusion criteria in Supplementary Fig. 2 could be used to reconstruct the focal-plane PT response. The $\varepsilon_A(x)$ and $\varepsilon_\varphi(x)$ calibration curves were then obtained from the measured cumulative and focal-plane PT responses. In the actual sample with probe beads, the calibrated focal-plane PT response (Supplementary Fig. 6, red) obtained from the measured cumulative PT response (Supplementary Fig. 6, blue) using the $\varepsilon_A(x)$ and $\varepsilon_\varphi(x)$ calibration curves was well overlapped with the direct measurements of the focal-plane PT response from the intrinsic OCT scattering signal of the collagen matrix (Supplementary Fig. 6, green). This confirms the validity of the PT calibration procedure in collagen matrices. The lateral $A_{\mathrm{PT}}(x, z_{\mathrm{f}})$ and $\varphi_{\mathrm{PT}}(x, z_{\mathrm{f}})$ as well as the axial $A_{\mathrm{PT}}(x_0, z)$ and $\varphi_{\mathrm{PT}}(x_0, z)$ profiles were obtained from the curve fit to both the direct-measurement and calibrated PT responses (Supplementary Fig. 6, black curves).

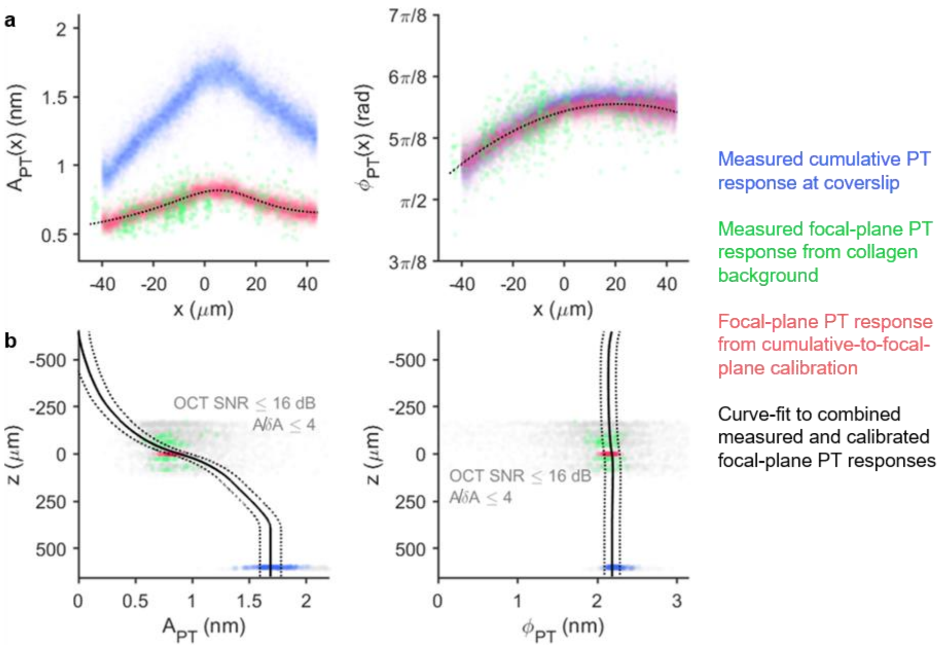


**Supplementary Fig. 6 Cumulative-to-focal-plane PT response calibration in collagen matrices. a**, Lateral profiles of measured cumulative (blue), measured focal-plane (green), and calibrated focal-plane from cumulative-to-focal-plane calibration (red), PT response amplitude (left), and phase (right) in the C3 sample. Black dotted curves represent Gaussian and quadratic fit to the combined focal-plane (i.e., red+green) $A_{\mathrm{PT}}$ and $\varphi_{\mathrm{PT}}$, respectively. The calibrated focal-plane PT response is in good agreement with the direct measurements available from the collagen matrix. **b**, The same comparison as in **a** but for the depth-dependent (axial) profiles at the centre of the light-sheet. Solid and dotted curves represent the best fit and ±1 median absolute difference between the best fit and the measurements, respectively.

**Cell-seeded fibrin constructs for live-cell imaging.** The fibrin construct generated intrinsic OCT scattering signal similar to collagen matrices, but the additional challenge for the purpose of live-cell imaging studies was the absence of the coverslip for cumulative PT response. This was necessary because it was desirable for cells to be exposed to the media and the environmental control of the incubation chamber, without the cell-seeded construct being sealed by the coverslip placed on top of the sample well. In other words, the only measurements of the PT response available in the live-cell imaging studies were from the intrinsic OCT scattering signal from the fibrin construct at the focal plane. Thus, no cumulative-to-focal-plane calibration was implemented in these studies. Instead, the direct measurements at the focal plane were supplemented by the measured focal-plane PT response in the "PT calibration sample" itself. The "PT calibration sample" in this case was another replicate of the same fibrin construct but containing no probe beads, no cells, and with coverslip placed on top of the sample well for cumulative PT response. Similar to the "PT calibration sample" for the collagen matrices, all available OCT scattering signal at the focal plane that passed the exclusion criteria in Supplementary Fig. 2 could be used to reconstruct the focal-plane PT response. Although the cumulative PT response from the coverslip was not necessary to characterize the lateral profile of the focal-plane PT response in this case, it was still needed to obtain the depth-dependent PT response at the centre of the light-sheet. The focal-plane PT response measured in the "PT calibration sample" (Supplementary Fig. 7, blue) was well overlapped with the direct measurements at the focal plane of the actual cell-seeded sample (Supplementary Fig. 7, green), which confirms the validity of the PT calibration procedure in cell-seeded fibrin constructs. The lateral $A_{\mathrm{PT}}(x, z_{\mathrm{f}})$ and $\varphi_{\mathrm{PT}}(x, z_{\mathrm{f}})$ as well as the axial $A_{\mathrm{PT}}(x_0, z)$ and $\varphi_{\mathrm{PT}}(x_0, z)$ profiles were obtained from the curve fit to the PT responses measured in both the actual cell-seeded and the "PT calibration sample" (Supplementary Fig. 6, black curves).

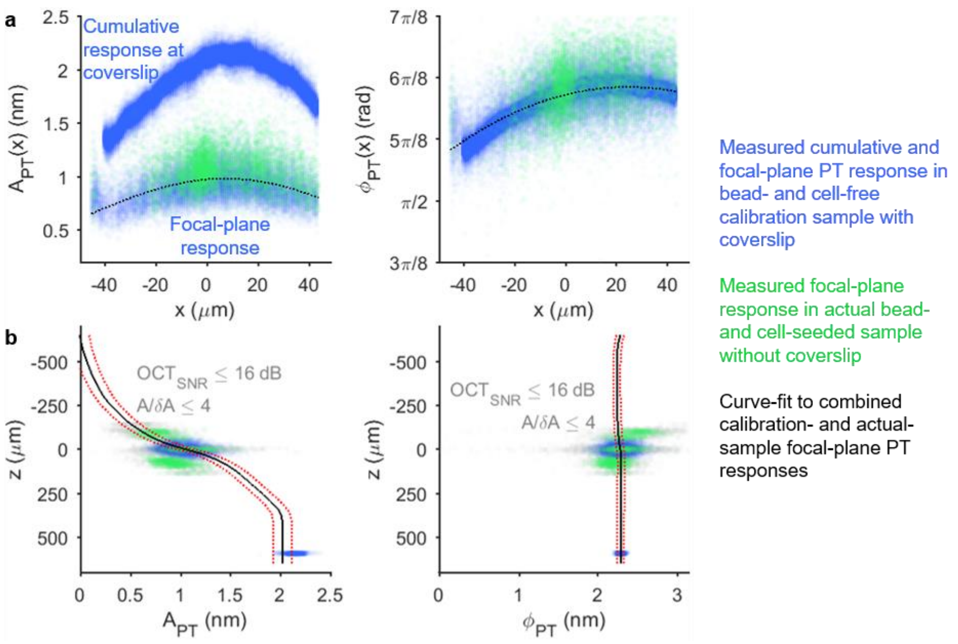


**Supplementary Fig. 7 PT response calibration in cell-seeded fibrin constructs for live-cell imaging. a**, Lateral profiles of PT response amplitude (left) and phase (right) measured in the actual cell-seeded fibrin construct (green) and the cell-free “PT calibration sample” (blue). Black dotted curves represent Gaussian and quadratic fit to the combined focal-plane (i.e., blue+green) $A_{PT}$ and $\varphi_{PT}$, respectively. The focal-plane PT responses measured in both samples are in good agreement with each other. **b**, The same comparison as in **a** but for the depth-dependent (axial) profiles at the centre of the light-sheet. Solid and dotted curves represent the best fit and ±1 median absolute difference between the best fit and the measurements, respectively.

### Supplementary Note 4: Light-sheet radiation-pressure force measurement

The radiation-pressure force from the light-sheet varies as a function of both $x$ and $z$ due to the Gaussian intensity profile of the light-sheet along its long axis. The light-sheet radiation-pressure force profile, $F_{\mathrm{rad}}(x,z)$, was measured by monitoring axial trajectories of beads in a viscous fluid under constant light-sheet beam power, according to a previously described depth-resolved method[6]. However, modifications were made to extend the measurement across $x$ to obtain the lateral force profile. Here, propulsion of beads was monitored in real-time via BM-mode imaging (as opposed to M-mode in Ref.[6]), with the fast-axis $x$ direction parallel to the long-axis of the light-sheet, and bead trajectory over time was analysed across the BM-mode frames (as opposed to M-mode A-scans in Ref.[6]). BM-mode frame rate of 76 Hz was sufficient for capturing bead acceleration by light-sheet radiation-pressure force.

Space-domain BM-mode OCT image was reconstructed with computational defocus correction along $x$ as described in Supplementary Note 2. Then, the BM-mode dataset was divided into eleven 9-μm wide lateral segments along $x$. Maximum intensity projection along $x$ was computed in each segment to produce eleven "M-mode-equivalent" bead trajectory images. Depth-resolved force reconstruction method[6] was performed on each image to produce the axial force profile in each lateral segment. Briefly, the instantaneous axial bead velocity, $\dot{z}(t)$, and acceleration, $\ddot{z}(t)$, were computed from the bead trajectory image. Then, instantaneous radiation-pressure force, $F_{\mathrm{rad}}(t)$, was computed by solving the axial equation of motion:

$$\frac{4}{3}\pi a^3 \rho_{\mathrm{bead}} \ddot{z}(t) = F_{\mathrm{rad}}(t) + \frac{4}{3}\pi a^3 g\left(\rho_{\mathrm{bead}} - \rho_{\mathrm{med}}\right) - 6\pi a \eta_{\mathrm{med}} \dot{z}(t), \tag{S16}$$

where $a$, $\rho_{\mathrm{bead}}$, $\rho_{\mathrm{med}}$, and $\eta_{\mathrm{bead}}$ denote the radius and mass density of the beads and the mass density and viscosity of the viscous fluid medium, respectively.

Due to the large number of bead trajectories to track in all eleven segments, an automated Radon transform-based bead trajectory tracking algorithm (see Supplementary Note 5 for details) was implemented instead of the previous manual tracking[6]. First, the axial force profiles from the 6$^{\mathrm{th}}$ and 7$^{\mathrm{th}}$ segments (two positions about the lateral centre of the light-sheet) were averaged to obtain $F_{\mathrm{rad}}(x_0,z)$. Then, the force magnitudes at the focal plane of the light-sheet from all lateral segments (i.e., force maxima along the axial profiles) were fit to a Gaussian curve as a function of $x$ to obtain $F_{\mathrm{rad}}(x,z_0)$. The 2D $F_{\mathrm{rad}}(x,z)$ profile was obtained via:

$$F_{\mathrm{rad}}(x,z) = F_{\mathrm{rad}}(x,z_0)\left[F_{\mathrm{rad}}(x_0,z) / F_{\mathrm{rad}}(x_0,z_0)\right]. \tag{S17}$$

However, the caveat for implementing the automated Radon transform-based bead trajectory tracking algorithm is that it underestimates the instantaneous axial velocity of the beads, resulting an underestimated reconstructed force magnitude (see Supplementary Note 5). To correct for this underestimation, the manual trajectory tracking procedure as implemented in Ref. 6 was performed only on the 6$^{\mathrm{th}}$ and 7$^{\mathrm{th}}$ segments, the results of which were averaged to obtain the "actual" peak force magnitude. Lastly, the $F_{\mathrm{rad}}(x,z)$ obtained from Eq. (S14) was multiplied by a correction factor of $F_{\mathrm{rad,manual}}(x_0,z_0) / F_{\mathrm{rad,automated}}(x_0,z_0)$.

### Supplementary Note 5: Automated algorithm for depth-resolved force reconstruction based on Radon transform

The previously described depth-resolved radiation-pressure force, $F_{\mathrm{rad}}(z)$, measurement method[6] is based on a semi-automated tracking of axial bead position over time, $z(t)$, using a coarse manual bead trajectory tracking to guide the search range of a subsequent automated algorithm. The instantaneous axial bead velocity, $\dot{z}(t)$, and acceleration, $\ddot{z}(t)$, are then obtained from the numerical derivatives of $z(t)$. However, the coarse manual bead trajectory tracking to map the 2D $F_{\mathrm{rad}}(x,z)$ profile for LS-pfOCE would involve manually tracking the trajectory of 200-400 beads each time a force measurement is needed (including both for $G^*$ reconstruction and routine system alignment check prior to each experiment), which is extremely tedious and time-consuming. Inspired by a previous method for blood flow velocity calculation from space-time image with Radon transform[7], an automated algorithm to extract depth-resolved axial bead velocity and acceleration was implemented using Radon transform over a sliding axial window (Supplementary Fig. 8).

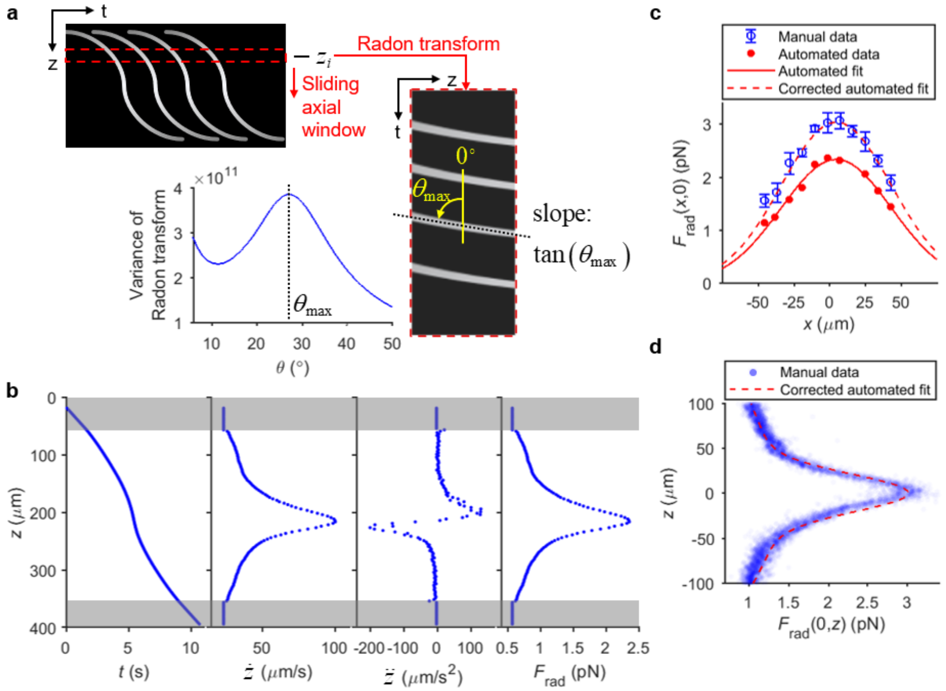


**Supplementary Fig. 8 Radon transform-based automated LS-pfOCE radiation-pressure force measurement.** **a**, Cartoon illustration of depth-resolved axial bead velocity (given by $\mathrm{slope} = \tan\theta_{\max}$) estimation from space-time bead trajectory image with Radon transform over a sliding axial window. **b**, Depth-resolved axial bead trajectory, velocity, acceleration, and $F_{\mathrm{rad}}$ magnitude obtained from Radon-transform-based reconstruction. **c**, Lateral profile of light-sheet $F_{\mathrm{rad}}$ obtained automated Radon-transform-based method before and after scaling correction, compared to results from manual-tracking-based method[6]. **d**, Axial profile of light-sheet $F_{\mathrm{rad}}$.

The automated algorithm is based on determining the “slope” of the trajectory of the bead within a small axial window centred at depth $z_i$ of the M-mode-equivalent (space-time) bead trajectory image using Radon transform (Supplementary Fig. 8a). The size of the axial window implemented here is 15 pixels. The angle made by the trajectory of the bead on the space-time image corresponds to the angle, $\theta_{\max}$, at which the variance of Radon transform magnitude is maximum. The average axial bead velocity within the $i^{\mathrm{th}}$ axial window, $\dot{z}(z_i)$, is given by:

$$\dot{z}(z_i) = (\delta z \cdot f_s)\tan(\theta_{\max}(z_i)), \tag{S18}$$

where $\delta z$ and $f_s$ denote the size of an axial pixel and the frame rate of the BM-mode acquisition, respectively. Then, the average axial bead acceleration within the $i^{\text{th}}$ axial window, $\ddot{z}(z_i)$, is estimated from rate of change of $\dot{z}(z_i)$ as a function of $z$ via:

$$\ddot{z}(z_i) = \dot{z}(z_i)\left[\left(\dot{z}(z_{i+1}) - \dot{z}(z_i)\right)/\delta z\right], \tag{S19}$$

where $z_{i+1} - z_i = \delta z$ for adjacent axial windows. Similarly, the axial bead trajectory $z(t)$ can be estimated from the cumulative summation of $\dot{z}(z_i)$. Lastly, the depth-resolved $F_{\text{rad}}(z_i)$ is computed from reconstructed $z_i$, $\dot{z}(z_i)$, and $\ddot{z}(z_i)$ as described in Ref. 6 (Supplementary Fig. 8b).

Due to the depth averaging over 15 axial pixels (sliding window size), the Radon-transform-based method underestimates the axial velocity $\dot{z}(z_i)$ compared to $\dot{z}(t)$ obtained from the numerical derivative of the manually tracked bead trajectory. This results in an overall underestimation of $F_{\text{rad}}$ magnitude, which can be corrected for with a scaling factor (Supplementary Fig. 8c, d). The scaling factor is determined by the ratio of the peak force magnitude, $F_{\text{rad}}(x_0, z_0)$, obtained from the manual-tracking-based method[6] to the automated Radon-transform-based method.

### Supplementary Note 6: Theoretical simulation of light-sheet radiation pressure

In our case, the radius of scattering particles is close to the wavelength of laser excitation, thus, we considered the Mie scattering mechanism. The simulation was conducted based on the assumptions that the light-sheet was launched along the $z$-axis and interacted with homogeneous and isotropic non-absorbing microspheres surrounded with non-absorbing medium, under the conditions of single scattering. The values of required parameters were based on our experimental practice. The beam shape coefficients (BSCs) were calculated based on Generalized Lorentz-Mie Theory (GLMT) by use of localized approximation[8]. The usual scattering coefficients of Lorentz-Mie theory depending on the properties of microparticles (size and refractive index) and the wavelength of laser beam were obtained from an open source algorithm[9]. The radiation pressure cross-section $C_{\mathrm{pr},z}(\mathbf{r})$ was computed via a finite series numerical integral over the calculated BSCs and Mie coefficients[10]. The photonic force was then calculated based on $F_{\mathrm{rad}} = \frac{2P_0}{cA} C_{\mathrm{pr},z}(\mathbf{r})$, where $P_0$ is the optical power, $c$ is the light speed in vacuum and $A$ is the cross-section area of the light-sheet. Assuming an elliptical-shaped Gaussian beam with $1/e^2$ beam radius along $x$ and $y$ directions of $w_{0,x}$ and $w_{0,y}$, respectively, we simply calculated $A = \pi w_{0,x} w_{0,y}$.

### Supplementary Data 1: Control experiments for live-cell imaging study

Two control experiments are performed to corroborate LS-pfOCE live-cell imaging results in Fig. 3. First, we test the effect of CytoD treatment on blank gels (i.e., fibrin constructs with no cell seeded), in order to verify that the difference in micromechanical properties observed in the normal versus CytoD conditions (Fig. 3b, e) is mediated by cellular activity. As expected, the stiffness of cell-free fibrin constructs is unchanged after the addition of CytoD (Supplementary Fig. 9a), indicating that the addition of the CytoD to the fibrin matrix does not alter its mechanical properties. Secondly, we test the effect of adding 50 μL of pure DMSO (which CytoD was dissolved in for the treatment condition) on the cell-mediated pericellular matrix stiffening, in order to verify that the difference in spatial variations of $G'$ and $R$ of the ECM around cells observed in the normal versus CytoD condition (Fig. 3f) is mediated by CytoD inhibition of actin polymerization. As expected, the variations in $G'$ and $R$ as a function of distance to cell follow the same trend before (Supplementary Fig 9b, red) and after (Supplementary Fig 9b, blue) the addition of DMSO. Notably, the exponential decay constants for the pericellular stiffness (inset of Supplementary Fig. 9b) observed both before and after the addition of DMSO here are consistent with that of the normal condition in Fig. 3f.

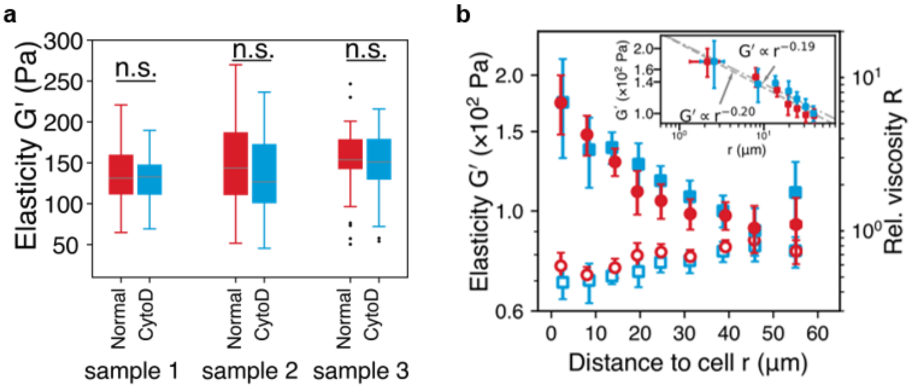


**Supplementary Fig. 9 Control experiments for live-cell imaging study. a**, LS-pfOCE $G'$ measured in blank (no cells seeded) fibrin constructs under normal condition (red) and after CytoD treatment (blue). n.s. indicates non-significant difference between group means based on Student's *t*-test. **b**, 3D LS-pfOCE $G'$ (left) and $R$ (right) in cell-seeded fibrin constructs as a function of distance to cell under normal condition (red) and after addition of DMSO (blue). Inset shows power law fits over the domain $r \leq 30$ μm. The decay exponent has a 95% confidence interval of –0.20 ± 0.064 (red) and –0.19 ± 0.054 (blue). Each data point represents mean ± standard deviation from $N \geq 20$ beads.

**Supplementary Data 2: Normalized stiffness gradient in the pericellular matrix from 3D live-cell micromechanical imaging**

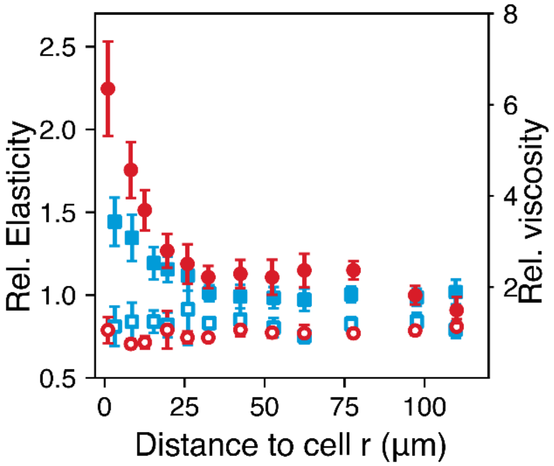


**Supplementary Fig. 10 Relative variations in pericellular viscoelasticity as a function of distance to cell.** 3D LS-pfOCE measurements in Fig. 3f normalized by the corresponding background (faraway regions) ECM mechanical properties of each data volume. The reduced stiffness gradient in the pericellular space can be clearly seen under the CytoD (solid blue) versus normal (solid red) condition.

### Supplementary Data 3: Comparison of cell-modified ECM micromechanical properties in the intermediate regions

3D LS-pfOCE measurements around normal versus CytoD-treated cells reveal a small increase in *G′* after the CytoD treatment in the intermediate ECM regions slightly further away from the cell (≥30 μm to cell in Fig. 3f). The comparison of *G′* and *R* in these intermediate ECM regions between the normal and CytoD conditions is provided in Supplementary Fig. 11. As discussed in the main body of the paper, we hypothesize that this mild stiffening after CytoD treatment in the intermediate ECM is related to the redistribution of the fibrin matrix from the pericellular space to the intermediate regions following the inhibition of cellular contractility.

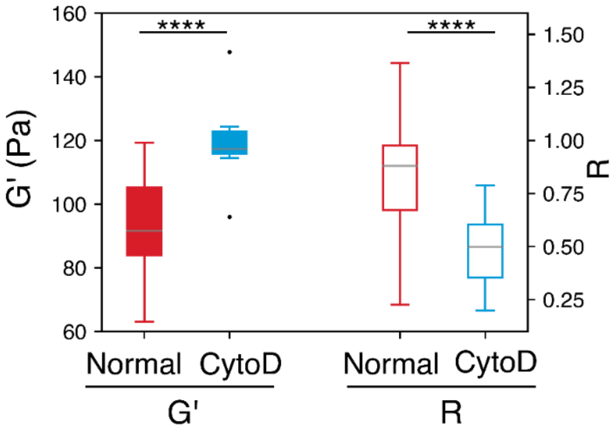


**Supplementary Fig. 11 Micromechanical properties of cell-modified fibrin ECM in the intermediate regions around cell.** Boxplots of *G′* (left) and *R* (right) measurements in Fig. 3f in the intermediate ECM regions ($r$ ≥30 μm) under normal (red) and CytoD (blue) condition. The background was calculated as the mean value of measurement micromechanics at regions away from cell body. Significant difference between group means based on Welch's *t*-test is indicated by ****$p$-value < 0.0001.

### Supplementary Data 4: Correlations between changes in micromechanical properties and local matrix deformations during time-lapsed monitoring of Cytochalasin D treatment

For dynamic monitoring of the cellular response to CytoD inhibitor, the cumulative changes in LS-pfOCE measurements were correlated to the signed cumulative displacement of each probe bead (Supplementary Fig. 12). The cumulative changes in $G'$ and $R$ at the $n^{\text{th}}$ time point were computed from $\Delta G'(t_n) = \sum_{i=2}^{n} G'(t_i) - G'(t_{i-1})$ and $\Delta R(t_n) = \sum_{i=2}^{n} R(t_i) - R(t_{i-1})$, while the signed cumulative bead displacement $S(t_n)$ was computed as described in Methods. Here, $S(t_n)$ represents a measure of local matrix deformation over time. The magnitude of $S(t_n)$ represents the magnitude of the bead displacement vector $\Delta\mathbf{r}_{\text{c}} = (\Delta x_{\text{c}}, \Delta y_{\text{c}}, \Delta z_{\text{c}})$ (Eq. (3) in Methods), while its sign represents the direction of the bead displacement along the major displacement axis (i.e., sign of $\Delta x_{\text{c}}$, $\Delta y_{\text{c}}$, or $\Delta z_{\text{c}}$, whichever is the largest across all beads) (Eq. (4) in Methods). Pearson linear correlation coefficients and $p$-values of $\Delta G'$ and $\Delta R$ to the cumulative bead displacement $S$ are indicated on Supplementary Fig. 10. As noted in the main body of the paper, an interesting observation (that has never been reported by other techniques to our knowledge) is that these changes occur in both the positive and negative directions, where an increased in $G'$ over time tends to coincide with a negative bead displacement (top right quadrant of Supplementary Fig. 12a) while a decrease in $G'$ over time tend to coincide with a negative bead displacement (bottom left quadrant of Supplementary Fig. 12a). Furthermore, a decreasing trend in $G'$ also tend to coincide with a higher initial stiffness. The temporal dynamics of each off the three bead displacement groups (blue, grey, and red shaded areas in Supplementary Fig. 12), representing three types of matrix deformation, are displayed as boxplots as a function of time in Fig. 3g–i.

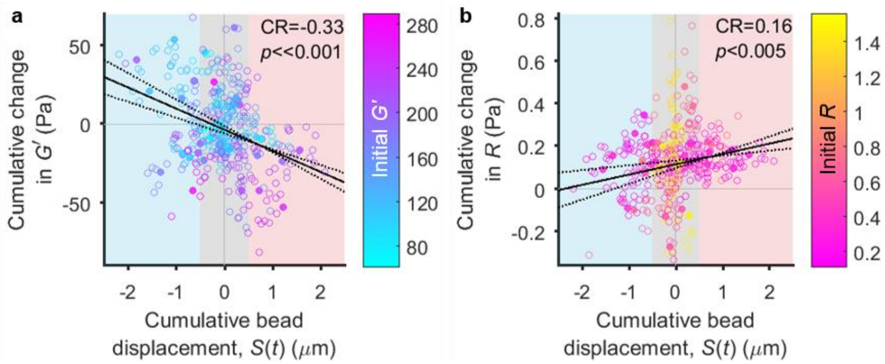


**Supplementary Fig. 12 Correlations between changes in micromechanical properties and local matrix deformations during time-lapsed monitoring of cytochalasin D treatment.** Cumulative change in **a**, $G'$ and **b**, $R$ as a function of cumulative bead displacement $S$ during time-lapsed LS-pfOCE measurements immediately after treatment with CytoD (see Methods for the calculation of $S$). $N = 34$ beads from 5 cells; each bead is color-coded by its initial $G'$ and $R$ values. All beads initially start at the origin and progress toward the solid marker at the last time point. Black solid and dotted lines represent linear fits and 95% confidence intervals, respectively. CR: Pearson linear correlation coefficient, $p$: $p$-value of Pearson linear correlation.

### Supplementary Discussion 1: Hardware requirements for LS-pfOCE data acquisition

Acquisition speed of our current LS-pfOCE is mainly limited by the low line-scan rate of the spectrometer and low performance of the stage motorized actuator. In our current implementation, line-scan rate of the spectrometer is 60 kHz with a frame rate of 425 Hz, which correspond to a data acquisition time of 15 s per slow axis position. Due to the limited acceleration and deceleration rate of our motorized actuator, waiting 6 s at each slow axis location prior to data acquisition is necessary for phase-sensitive measurements. This amounts to a total data acquisition time of 227 min with 117 min of actual OCE image acquisition and an additional 50 min of wait time for the motorized stage actuation. With a state-of-the-art swept-source OCT capable of multi-MHz A-scan rate, incorporating a resonant scanner and high-fidelity motorized stepper[11], the total acquisition time for a 3D LS-pfOCE dataset could be reduced to <10 min when acquiring 512 B-scans (slow axis positions). However, we note that attention must be given to the phase stability of the system when implementing high-speed scanning in practice. Thus, a *practical* robust 3D LS-pfOCE data acquisition may be expected within a few tens of minutes, with faster scanning while ensuring adequate phase stability.

Data transfer rate (limited by hardware) is another important consideration when designing/implementing the LS-pfOCE system and experiments. For our current implementation with a 425-Hz frame rate, 120 A-scans per frame, and 2048 camera pixels, the required data transfer speed is ~156 MB/s with 12-bit precision, which sets the minimum required data transfer speed of the frame grabber. A spectrometer with two tap configuration is typically used to speed up data transfer in high-speed spectral-domain OCT, while data transfer rate may potentially be less of a concern for swept-source OCT. Moreover, large data storage drives are another prerequisite for LS-pfOCE systems. When conducting 3D volumetric imaging, ~1.5 TB of binary data was acquired for each full 3D LS-pfOCE volume (2048 camera pixels × 120 A-scans × 467 slow-axis position × 6400 temporal frames per slow axis position). Notably, these hardware considerations for implementing an LS-pfOCE system will highly depend on the application-specific image acquisition parameters (e.g., FOV, frame rate, number of temporal frames).
